# Structure dynamics of ApoA-I amyloidogenic variants in small HDL increase their ability to mediate cholesterol efflux

**DOI:** 10.1101/2020.04.08.031211

**Authors:** Oktawia Nilsson, Mikaela Lindvall, Laura Obici, Simon Ekström, Jens O. Lagerstedt, Rita Del Giudice

## Abstract

Specific mutations in Apolipoprotein A-I (ApoA-I) of high-density lipoprotein (HDL) are responsible for a late-onset systemic amyloidosis. Carriers do not exhibit increased cardiovascular disease risk despite reduced levels of ApoA-I/ HDL-cholesterol. To explain this paradox, we show that the HDL particle profile of L75P and L174S patients presents a higher relative abundance of the 8.4 nm vs 9.6 nm particles, and that serum from patients, as well as reconstituted 8.4 and 9.6 nm HDL particles (rHDL), possess increased capacity to catalyze cholesterol efflux from macrophages. Synchrotron radiation circular dichroism and hydrogen-deuterium exchange revealed that the variants in 8.4 nm rHDL have altered secondary structure composition and display a more flexible binding to lipids compared to their native counterpart. The reduced HDL-cholesterol levels of patients carrying ApoA-I amyloidogenic variants are thus balanced by higher proportion of small, dense HDL particles and better cholesterol efflux due to altered, region-specific protein structure dynamics.

## Main

Amyloidoses are a broad group of diseases caused by protein instability and misfolding that leads to pathological aggregation of proteins as amyloid deposits in tissues and organs^1^. These amyloid deposits are either localized, as in Alzheimer’s and Parkinson’s diseases, or systemic, as is the case of transthyretin and light-chain amyloidoses. The different types of amyloidoses commonly lead to severe and age-related damage of the tissues where aggregation takes place. The understanding of the determinants leading to protein fibril formation and amyloidosis development is thus key for finding ways to develop efficient treatments for the affected individuals.

Apolipoprotein A-I (ApoA-I), associated with hereditary systemic amyloidosis, has a key role for the transportation of cholesterol and lipids in the circulation between peripheral tissues and the liver^2^. Following its synthesis in the liver and intestines, ApoA-I mediates this transportation by the formation of high-density lipoprotein (HDL) particles. These load cholesterol and lipids from macrophages at the artery wall via a regulated efflux mechanism catalyzed by the cellular ATP-binding cassette (ABC)A1 receptor. HDLs are then carried to the liver in the bloodstream for cholesterol processing and/or excretion as bile. This is an important process since well-balanced levels of cholesterol, particularly at the artery wall, may reduce the build-up of atherosclerotic plaques and hence the risk of cardiovascular disease (CVD). HDL particles are further modified by interaction with additional cellular receptors (ABCG1 and scavenger receptor BI (SR-BI)) and with soluble enzymes, including the lecithin-cholesteryl acyl transferase (LCAT) protein^3,4^. During these processes the HDL particle grows from the initially formed discoidal pre-beta HDL species to larger, spherical HDL particles, a process that is facilitated by flexibility in the ApoA-I structure^5^. The interaction with these receptors and soluble enzymes, as well as the protein flexibility, are critical for HDL biogenesis. The qualitative features and the size-distribution of the HDL species, rather than the absolute quantity of ApoA-I/HDL, are indeed regarded as major determinants for function^6^ and for the prevention of CVD^7^.

A high degree of flexibility in the ApoA-I structure is thus important for lipid-binding and functionality of ApoA-I in HDL. However, in ApoA-I variants carrying single amino acid substitutions, this flexibility might be causative for increased susceptibility to proteolytic remodeling and subsequent protein aggregation. Indeed, this is the case for twenty-three currently known human amyloidogenic variants of the *APOA1* gene that lead to progressive accumulation of ApoA-I protein in liver, heart, kidneys, larynx, skin and/or testis^8–12^. These mutations are localized to two major regions of the ApoA-I structure, either within residues 25 to 75 in the N-terminal domain or within residues 170 to 178 in the central domain of ApoA-I. The reason for the high occurrence of amyloid-prone variants in these two regions is not known, but this might indicate that these regions have specific functions in lipid-association and/or in protein structure dynamics^13^. Interestingly, carriers of several ApoA-I variants, including Gly26Arg ^14^, Leu75Pro^15,16^, and Leu174Ser^17,18^, have decreased blood levels of ApoA-I, yet do not have increased risk of CVD. Recently, *in vitro* cell studies using recombinant lipid-free and HDL-reconstituted ApoA-I amyloidogenic variants provided an explanation to this apparent paradox, as these variants showed a significantly higher cholesterol efflux capacity compared to the native protein^19^. However, the structural basis for the improved catalytic function of the ApoA-I variants is not understood. It is also not clear how the complexity and HDL-species distribution in circulation contribute to the improved efficacy of the variants. Here, we use clinical samples from Leu75Pro (L75P) and Leu174Ser (L174S) patients, as well as from matched control individuals, to investigate their unique HDL particle distribution and functionality. In order to explain the *in vivo* phenotype of the L75P and L174S variants, synchrotron-radiation circular dichroism (SRCD), hydrogen-deuterium exchange (HDX) mass-spectroscopy on reconstituted HDL (rHDL), and the measurement of cholesterol efflux from macrophages to both rHDL and patient serum samples are used to analyze the functionality and structure of discoidal pre-beta HDL particles of defined sizes.

## Results

### Patients carrying ApoA-I amyloidogenic variants have distinctive HDL patterns

Serum samples from patients carrying ApoA-I amyloidogenic variants (heterozygotic for either L75P or L174S) and from unrelated control subjects were depleted for Apolipoprotein B (potential cholesterol acceptor) prior to the analyses. To examine the levels and profile of HDL species, equal volumes of serum samples were separated by denaturant and native PAGE and analyzed by western blot for ApoA-I protein content by using anti-human ApoA-I antibodies. Serum from L75P heterozygotic patients presented lower amounts of total ApoA-I protein (33 % reduction compared to control subjects, Figure 1A), which agrees with previous reports^16,17,20^, whereas ApoA-I levels in serum of L174S patients were similar to those of control subjects. Analysis of the samples by native western blot supported this observation and also revealed differences in the distribution of HDL species (Figure 1B). Specifically, serum from control subjects displayed an additional species of particles around 12 nm and also a higher relative abundance of the 9.6 nm particles in relation to the 8.4 nm particles (9.6 nm to 8.4 nm ratio is 1.09) compared to serum from L75P and L174S patients (ratios of 0.57 and 0.71, respectively) (Figure 1B, right panel).

**Figure 1.**
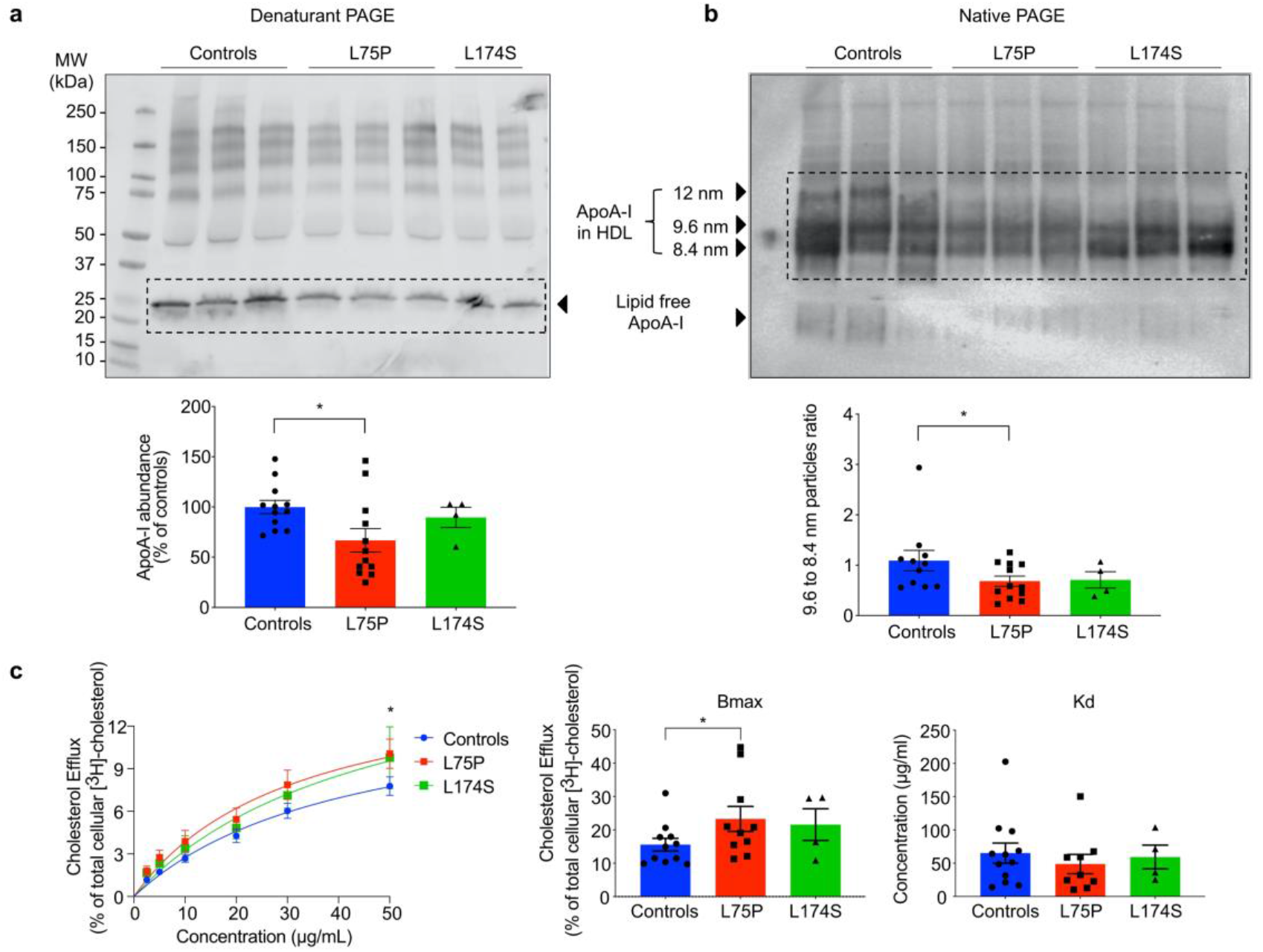
Quantitative and qualitative analysis of HDL from patients with ApoA-I amyloidosis. **a,** Patients carrying ApoA-I amyloidogenic variants possess reduced levels of plasma ApoA-I. Proteins from serum sample from patients and control subjects were separated by denaturant PAGE, analyzed by western blot by using anti-human ApoA-I antibody (upper panel) and ApoA-I levels were quantified by using ImageJ software (lower panel). **b,** Patients carrying ApoA-I amyloidogenic variants have a distinctive HDL pattern. Proteins from serum samples from patients and control subjects were separated by native PAGE and analyzed by western blot by using anti-human ApoA-I antibody (upper panel). The ratio between the abundance of 9.6 nm and 8.4 nm particles was calculated and is displayed in the lower panel. **c,** HDL from patients show improved ability to promote cholesterol mobilization. Cholesterol efflux experiments were performed by using serum samples as acceptors of cholesterol, in dose-response experiments. Radioactive cholesterol-loaded J774 macrophages were treated with serum samples, normalized for the amount of total ApoA-I, and incubated for 4 h. The experimental data (left panel) were fitted and Bmax (middle panel) and Kd (right panel) were calculated according to Michaelis - Menten equation. Data shown are the mean ± SEM and significance is calculated according to two-way ANOVA (c, left panel, *p<0.05), or one-way ANOVA (c, middle panel, *p<0.5. N = 11 for L75P and controls, and N = 4 for L174S donors. Representative samples from three donors per group are shown in panels a and b.

### HDL from patients carrying ApoA-I amyloidogenic variants show improved ability to promote cholesterol mobilization

Reconstituted 9.6 nm HDL particles containing ApoA-I variants have been shown to be more efficient than the wild-type (WT) counterpart at mediating cholesterol efflux from macrophages^19^. In order to test if HDL from patients carrying L75P and L174S amyloidogenic variants possess an improved ability to mediate cholesterol mobilization, dose-response cholesterol efflux experiments were performed by using serum samples as acceptors of cholesterol. Cholesterol-loaded J774 macrophages were incubated (4 h) with serum samples from L75P and L174S patients, or from control subjects, in a dose-dependent manner normalized for the amount of total ApoA-I. Spontaneous cholesterol release into the media was measured and the efflux for each treatment subtracted for this value.

As shown in Figure 1c, patients with the L75P substitution displayed a 33% increase in cholesterol binding capacity (Bmax) as compared to control subjects (p<0.05) but with no difference in binding affinity (Kd). A similar increase in the efflux capacity of serum from L174S patients was observed. However, the increase in Bmax did not reach statistical significance, most likely due the lower number of clinical samples available for this variant. The improved efflux capacity observed for the amyloidogenic ApoA-I patients may be sex-specific as serum from male patients showed the larger difference in Bmax compared to male control subjects (Figure S1).

In order to evaluate if the relatively higher serum concentrations of 8.4 nm HDL particles (compared to the 9.6 nm HDL particles) contributed to the improved cholesterol mobilization, cholesterol efflux experiments were performed by using reconstituted HDL (rHDL; POPC:ApoA-I) particles with sizes of either 8.4 or 9.6 nm. The effect of unesterified free cholesterol (FC) already being present in the synthesized rHDL particles (POPC:FC:ApoA-I) was also evaluated.

In general, several of the rHDL particles containing the amyloidogenic variants displayed greater ability to promote cholesterol efflux compared to WT ApoA-I (Figure 2). In particular, 9.6 nm POPC:FC:ApoA-I particles containing either L75P or L174S ApoA-I protein were characterized by a higher cholesterol binding capacity (from 20 to 30 % higher cholesterol binding at all concentrations tested; Figure 2, top panels) when compared to WT rHDL. Similarly, it was previously observed that 9.6 nm POPC:ApoA-I rHDL containing either L75P or L174S amyloidogenic variant showed improved cholesterol efflux ability, however, in that case due to a higher affinity to cholesterol^19^. Interestingly, in the case of 8.4 nm rHDL, only the particles containing the L75P variant showed an increased cholesterol binding capacity. This increase was particularly pronounced for L75P rHDL particles synthesized without FC (POPC:ApoA-I), which were also characterized by a higher cholesterol binding affinity (lower Kd) compared to WT- and L174S-containing rHDL.

**Figure 2.**
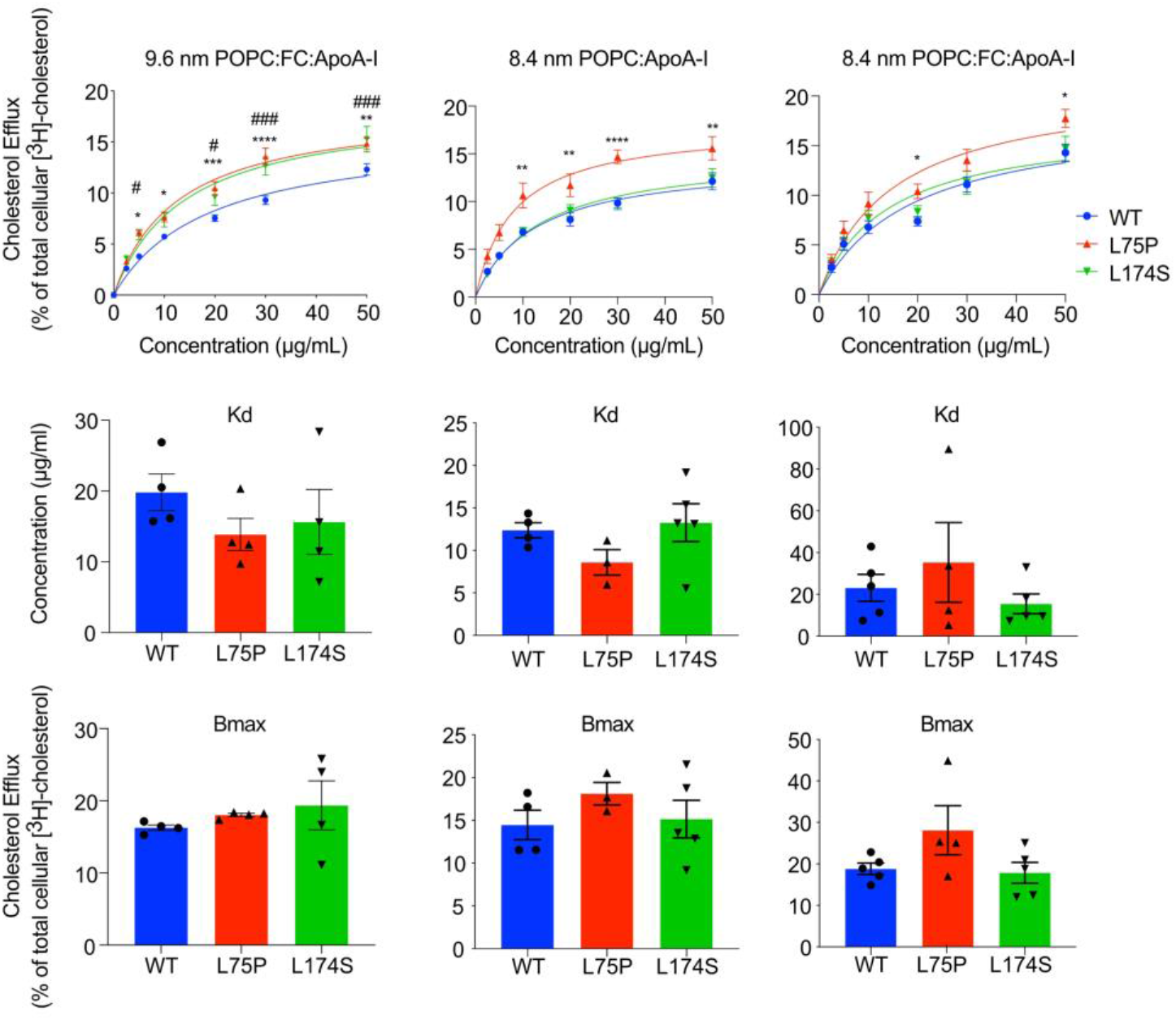
Cholesterol efflux ability of reconstituted HDL of different sizes. 9.6 and 8.4 nm reconstituted HDL (rHDL) were produced by incubating recombinant ApoA-I amyloidogenic variants with POPC, in the presence or absence of cholesterol, and tested for their ability to mediate cholesterol efflux from J744 macrophages, in dose-response experiments. The experimental data (upper panel) were fitted and Kd (middle panel) and Bmax (lower panel) were calculated according to Michaelis - Menten equation. Data shown are the mean ± SEM and significance is calculated according to two-way ANOVA (*p<0.05, **p<0.005, ***p<0.001, ****p<0.0001, and ## p<0.005, ### p<0.001 for L75P and L174S rHDL, respectively, as compared to WT rHDL). N = 4-5.

These observations suggest that the improved efflux ability of HDL from L75P patients (Fig 1C) is due to the higher levels of 8.4 nm particles compared to controls.

### ApoA-I amyloidogenic variants in 8.4 nm particles are characterized by a higher structural flexibility

Far-UV circular dichroism (CD) spectroscopy was used to analyze protein conformation and stability of the ApoA-I variants, as well as the WT protein, in 8.4 nm rHDL.

The estimation of the secondary structure of the proteins in rHDL, performed by synchrotron radiation CD, revealed interesting differences in the folding of the amyloidogenic variants and the native protein (Figure 3a). In the absence of FC (POPC:ApoA-I), the L75P variant in 8.4 nm rHDL particles showed a remarkably high content of α-helices (90 % for L75P compared to 64 % for WT), and a low content of turns and unordered structures. The L174S variant, instead, was characterized by a lower percentage of α-helical structure (39 %) accompanied by higher proportions of turns, beta-strands and unordered structures. In the presence of FC, the amount of α-helical structure of both WT and L174S in 8.4 nm rHDL were increased compared to the non-FC particles; however, the L174S α-helical content was still significantly lower than that for both WT and L75P.

**Figure 3.**
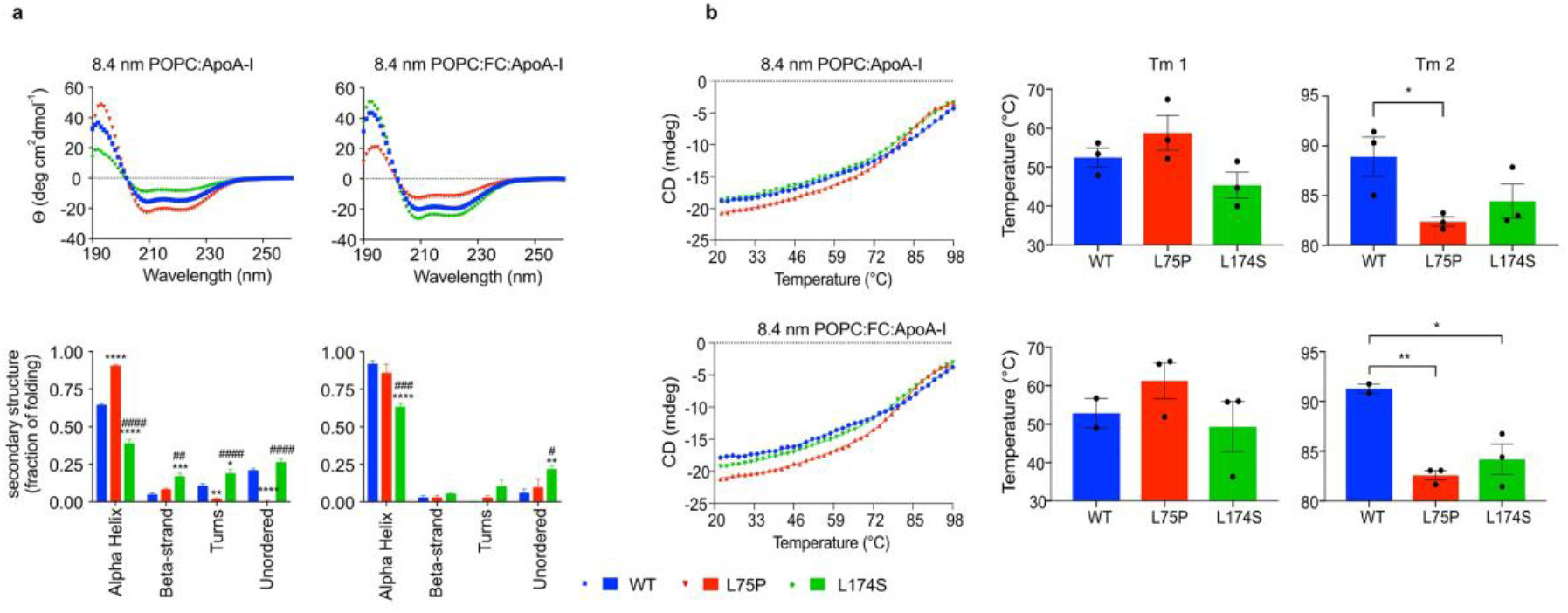
Conformation and stability of ApoA-I amyloidogenic variants in 8.4 nm rHDL. **a**, SRCD spectroscopy analysis of ApoA-I variants in 8.4 nm rHDL particles, in the presence or absence of FC. SRCD spectra (upper panels) were deconvoluted with the CONTINLL algorithm for the estimation of the ApoA-I proteins’ secondary structure (lower panels). **b**, thermal stability of ApoA-I variants in 8.4 nm rHDL particles, in the presence or absence of FC. The unfolding curves (left panels) were obtained by monitoring the CD signal amplitude at 222 nm as a function of temperature and Tm (right panels) were calculated by fitting the experimental data with a biphasic non-linear regression. Data shown are the mean ± SEM and significance is calculated according to (a) two-way ANOVA (*p<0.05, **p<0.005, ***p<0.001, ****p<0.0001, and ## p<0.005, ### p<0.001 #### p<0.0001 for L75P and L174S rHDL, respectively, as compared to WT rHDL, or (b) one-way ANOVA (*p<0.05, **p<0.005). Data shown are the mean ± SEM. N = 3.

ApoA-I proteins in 8.4 nm rHDL were next tested for their thermal stability by monitoring changes in the CD signal at 222 nm. As already reported for 9.6 nm POPC:ApoA-I particles^11^, thermal denaturation of the rHDL particles follows a biphasic unfolding process, showing two transition temperatures (Tm). These are considered to reflect structural rearrangement of the protein (Tm1) and dissociation of protein from lipids (Tm2). As shown in Figure 3b, the two ApoA-I variants showed similar Tm1 when compared to the WT (middle panels) but were both characterized by lower Tm2 (right panels), suggesting a higher protein flexibility and/or a lower affinity for lipids. Of note, the thermal denaturation profiles appeared to be unaffected by the presence of FC, suggesting that cholesterol does not contribute to the thermal stability of neither the ApoA-I protein nor to the protein-lipid association in rHDL. To further explore the thermal denaturation process and the species formed, 8.4 nm POPC:ApoA-I rHDL were incubated at 20°C, 65°C (temperature above Tm1), 85°C (slightly above Tm2 for the variants but below Tm2 of the WT) and 98°C (temperature at the end of the thermal unfolding process and clearly above Tm2 for all three samples). The collected samples were then separated by native PAGE and analyzed by western blot (Figure 4). Both ApoA-I variants were characterized by higher amounts of the lipid-free species at equilibrium (20°C; CTRL), which was particularly pronounced in the case of the L75P variant (5.3-fold higher level of lipid-free L75P protein compared to WT). The relative abundances of lipid bound and lipid-free species remained unchanged upon heating at 65°C. At 85°C, the L75P sample showed the highest amount of lipid-free species, followed by the L174S sample, in agreement with the observed lower Tm2 (Figure 3b). At the end of the thermal denaturation process (98°C), almost all of the L75P and L174S proteins were in their lipid-free states whereas a significant amount of the WT protein (18%) was still associated with lipids. The more pronounced dissociation of the variants from lipids and the lower temperature at which this occurred indicate that the ApoA-I variants in rHDL are characterized by a much higher flexibility compared to the WT protein.

**Figure 4.**
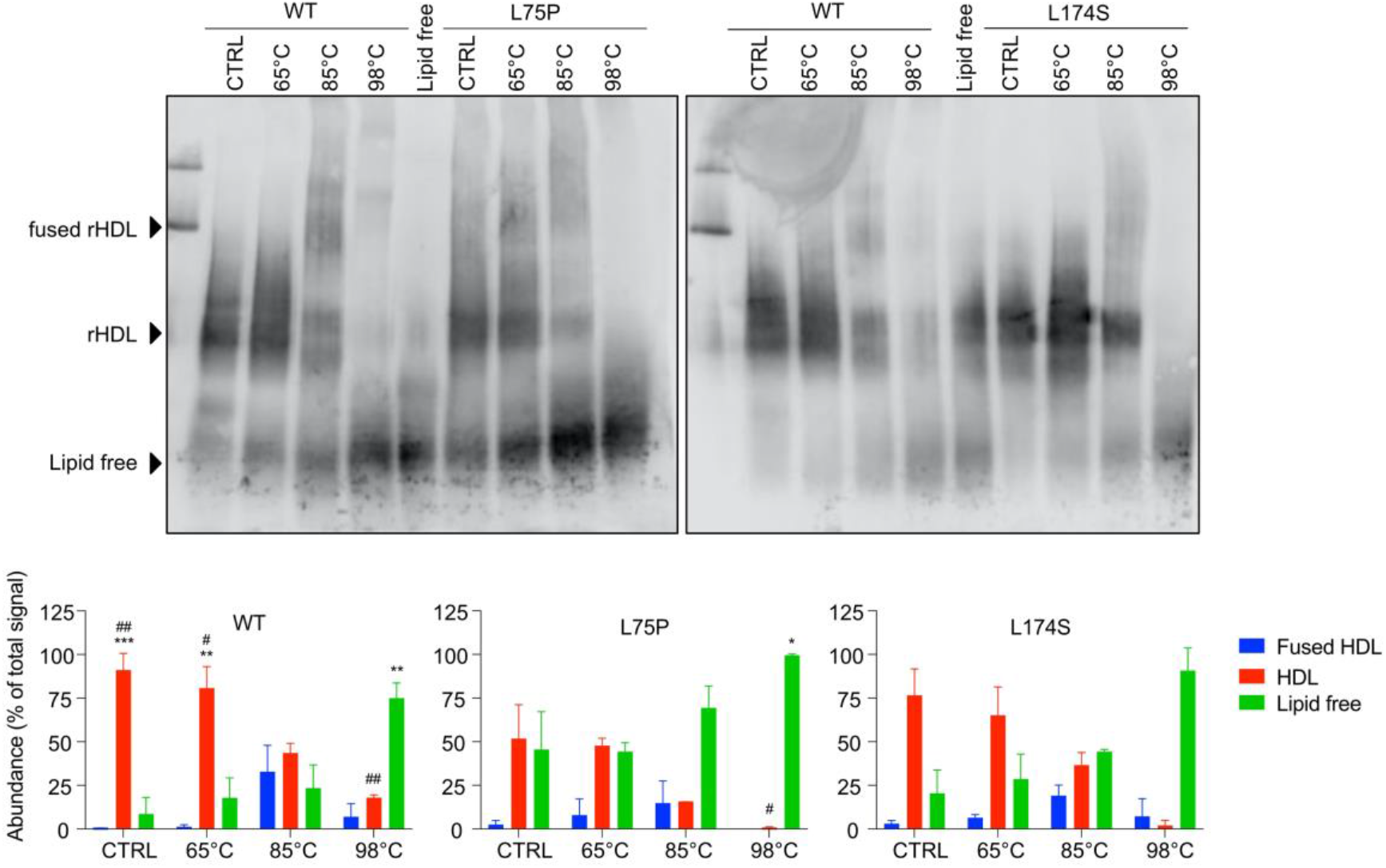
Temperature-induced unfolding and lipid-dissociation of ApoA-I amyloidogenic variants in 8.4 nm rHDL. 8.4 nm rHDL particles, in the absence of FC, were incubated at 20°C (CTRL), 65°C, 85°C and 98°C, separated by native PAGE and analyzed by western blot by using anti-human ApoA-I antibody (upper panels). Lipid-free ApoA-I was used as control. The species obtained upon thermal denaturation were quantified and the amount of each species was expressed as percentage with respect to the total signal (lower panels). Data shown are the mean ± SD and significance is calculated according to two-way ANOVA (*p<0.05, **p<0.005, ***p<0.001 for groups as shown respect to fused HDL, # p<0.05, ## p<0.005 for groups as show respect to lipid-free protein). N = 2 (L75P and L174S rHDL), or 4 (WT rHDL).

SRCD and thermal denaturation provide information on the global protein secondary structure, as well as on protein stability and flexibility but do not describe which protein regions or domains are specifically affected by the L75P and L174S substitutions. To address this, hydrogen-deuterium exchange mass spectrometry analysis (HDX-MS) was used to analyze domain-specific protein flexibility of the ApoA-I proteins in 8.4 nm rHDL. The differential heatmaps between the variants and the WT rHDL (Figure 5a-b) indicated an overall increase in deuterium uptake for both the ApoA-I amyloidogenic variants (heatmaps for the individual proteins are shown in Figure S2). Notably, both L75P and L174S variants showed region-specific structural flexibility, with the L75P variant presenting a stronger destabilizing effect in the regions close to the mutation site (50-73 and 74-105 regions), whereas the L174S variant was characterized by a decrease in stability in the 126-140 and 170-180 regions (deuterium uptake for the peptides covering the full proteins sequence are shown in Supplementary data file S1). Furthermore, the differential heatmap showing L75P vs L174S deuterium uptake (Figure 5c) depicted a much flexible N-terminal domain of L75P rHDL, particularly in the region 50-73, where no significant difference between the WT and L174S rHDL could be detected (Figure 6, peptide ID 35 and 40), and a higher exchange rate for the L174S in the 170-180 region already at the shortest deuterium exposure time (Figure 6, peptide ID 114).

**Figure 5.**
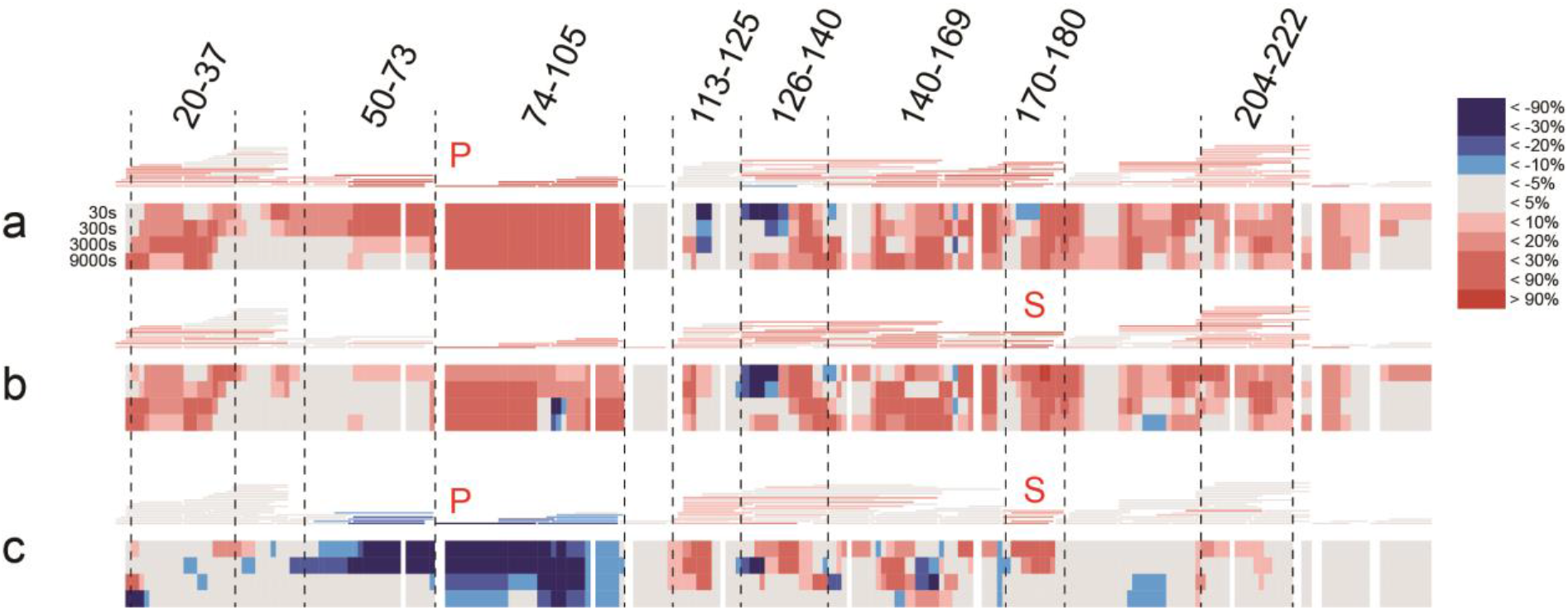
Hydrogen-deuterium exchange mass spectrometry reveals region with high flexibility in 8.4 nm rHDL. **a**, Differential HDX heatmap of L75P vs WT, **b**, L174S vs WT and **c**, L174S vs L75P proteins in 8.4 nm POPC particles. HDX peptide coverage is shown by the bars above each heatmap. The heatmaps show the deuterium uptake at the different time points (30 s, 300 s, 3000 s and 9000 s). Cold color = slower exchange; warm color = faster exchange.

**Figure 6.**
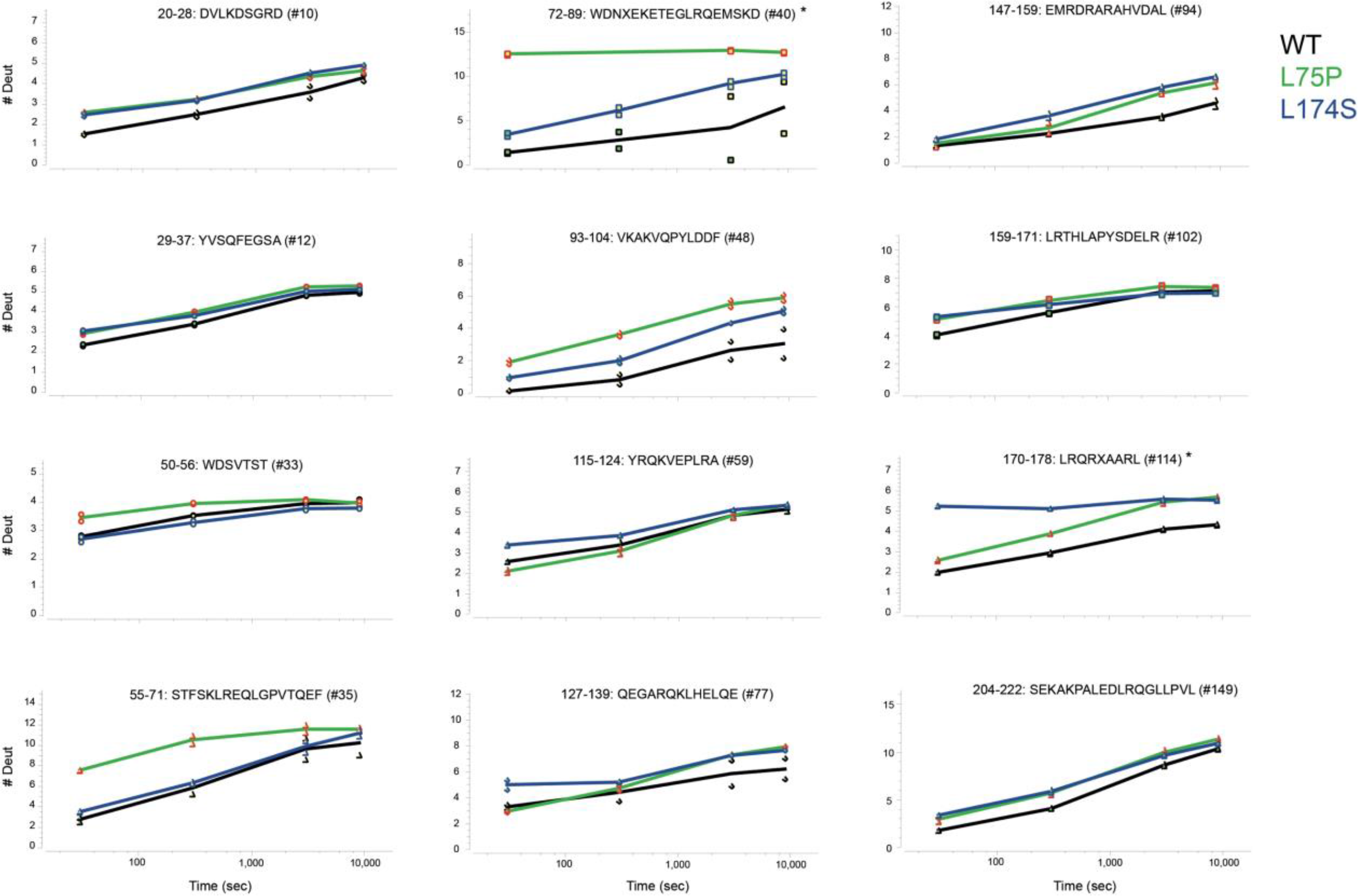
Deuterium uptake of selected ApoA-I peptides. Individual deuterium uptake plots for the ApoA-I were the aminoacidic substitutions led to a decreased structural stability compared to WT protein. The peptide ID number for the selected experiment and sequence are reported above each plot. Black trace (WT), green (L75P) and blue (L174S). * indicates peptides containing the aminoacidic substitution.

## Discussion

Subjects affected by ApoA-I hereditary systemic amyloidosis are characterized by low levels of serum ApoA-I and HDL; however, although they also present reduced levels of HDL-cholesterol, patients do not show higher risk of developing CVD^15–18^. We previously showed that ApoA-I amyloidogenic variants in reconstituted 9.6 nm HDL particles exhibit improved cholesterol efflux capability and we therefore hypothesized that this enhanced functionality could serve to compensate the unfavorable lipid profile of amyloidogenic ApoA-I carriers^19^. The present study translates these findings and verifies that the amyloidogenic ApoA-I variant L75P in serum from human patients displays improved cholesterol efflux activity (Fig 1c). A similar trend for the L174S variant was observed but was not statistically significant, possibly due to the low number of available serum samples from L174S patients.

This study also establishes that the ApoA-I amyloidogenic variants L75P and L174S present unique patterns in pre-beta HDL particle distribution in serum from the patients. In particular, the level of smaller HDL particles (8.4 nm as compared to 9.6 nm particles) is higher in L75P and L174S patients in comparison to the controls, and the larger 12 nm particles were relatively more abundant in the controls (Fig 1b). HDL particle size has been shown to be a critical determinant of ABCA1-mediated cholesterol efflux in macrophages, with small dense particles being the most efficient mediators^6,21^. The shift towards a larger proportion of small, dense particles in serum from patients could therefore potentially provide the explanation for the observed improvement in cholesterol efflux. However, a contributing role of the amino acid substitutions of the variants *per se* could not be ruled out. To investigate this possibility, we compared the functionality of small, dense and variant-specific rHDL particles with defined sizes and protein/lipid compositions. The fact that the cholesterol efflux to the 9.6 nm rHDL acceptor particles was higher for both variants compared to the WT ApoA-I rHDL particles clearly indicated that the substitution at either residue 75 or 174 have a direct impact on protein/rHDL functionality. Interestingly, when in 8.4 nm rHDL particles, only the L75P variant showed elevated capacity for cholesterol efflux, and this difference appeared to be particularly significant for particles reconstituted without free cholesterol. The data thus indicate that the L75P and L174S substitutions affect, at least partly, different molecular transitions/mechanisms and that this occurs in a particle-size-dependent manner. This may not be surprising, as ApoA-I is likely to adopt specific conformations in HDL particles with different sizes^5^. Similarly, since ApoA-I proteins showed differences in functionality between the cholesterol/non-cholesterol particles, ApoA-I may adopt specific conformations that also depend on the HDL lipid composition.

The organization of the primary structure into amphipathic alpha helices is a key feature in the lipid binding process of the ApoA-I protein. The L75P variant in 8.4 nm rHDL, without free cholesterol, was shown to have a significantly larger proportion of alpha-helices compared to both L174S and WT rHDL particles (Fig 3a, left column). The high alpha helical content of the L75P protein in 8.4 nm rHDL may thus indicate a readiness of the protein to accept cholesterol, and that preloading of the particles with cholesterol (Fig 3a, right column) triggers structural transitions to higher proportions of alpha helical secondary structure also in the L174S and WT proteins. Considering the central role of ApoA-I amphipathic alpha helices in HDL particle formation, the high level of beta-strand/turns and unordered structure in the L174S variant in 8.4 nm rHDL, in particular the cholesterol-free particles, is intriguing. How the characteristic structural organization of L174S in 8.4 nm HDL relates to protein function, and also to the amyloidogenic propensity of L174S protein, is not clear, but may partly explain the observed differences in cholesterol efflux capacity of the two variants (Fig 2; 8.4 nm rHDL), as well as the reduced phospholipid-binding capacity of lipid-free L174S^19^.

We also found that the overall stability of the variants in 8.4 nm rHDL particles is not affected (Tm1 in Fig 3b), although their interactions with lipids are weaker than that observed for the WT protein (Tm2 in Fig 3b). This finding indicates that the amyloidogenic amino acid substitutions impose regional changes in protein backbone flexibility without substantially affecting the structural elements that contribute keeping the integrity of the entire protein. These conclusions are further supported by the qualitative analysis of rHDL integrity, which shows that the variants, in particular the L75P variant, are more prone to dissociate from the phospholipids/cholesterol of the lipid-protein complexes.

Sequence analysis of the ApoA-I protein described one globular domain (residues 1-43) followed by consecutive alpha helices (h1 to h10) with lengths of 11 or 22 amino acids^22^. In the discoidal HDL particles, two ApoA-I molecules are organized in an antiparallel fashion (double belt) with a h5-h5 interaction between the two monomers^23^. The L75P and the L174S substitutions are in the centers of h2 and h7, respectively. The h2 and h7 helices from the two ApoA-I monomers in discoidal HDL particles are partially overlapping, which brings residues 75 and 174 in relatively close proximity in the HDL particles. Moreover, a proline at position 75 is likely to create a kink that effectively breaks the 22mer h2 helix into two shorter 11mer helices, i.e., the same length as h3 and h9.

HDX-MS has previously been successfully applied for the study of the ApoA-I dynamics in rHDL ^24–27^. In a recent publication, the authors performed HDX on both lipid-free and -bound forms of three ApoA-I variants (F71Y, L159R and L170P). Although these variants were studied in rHDL particles with a different size (11-12 nm for F71Y, L159R and L170P vs 8.4 nm for L75P and L174S) and lipid composition (DMPC vs POPC), and despite the fact that comparing HDX data is inherently complicated^28^, the present study describes common trends with the previous HDX data. Indeed, a general reduction in uptake when going from lipid-free to lipid-bound form was observed (Fig S2), as well as a decreased protection of the region spanning the h5 pair (L159R and L170P). Moreover, the HDX-MS analyses of 8.4 nm rHDL particles (Fig 5) show a specific increase in backbone flexibility in the 55 to 89 region for the L75P variant and in the 170 to 178 region for the L174S variant (Fig 6), and a general increase in solvent accessibility for the variants compared to the WT protein in 8.4 nm rHDL particles (Fig 5, Fig S3). Overall the variants provide a more dynamic protein-lipid interaction that appears to be beneficial for their function.

The HDX-MS analysis of L75P and L174S in rHDL showed that deuterium uptake increases in regions that are already relatively flexible in the WT protein. Uptake is greatly increased at the mutation site, but an increase in deuteration in the h5/5 region and a broadening of solvent exposure towards the 4/6 helices could also be observed.

It appears that substitutions that break or weaken the alpha-helix hydrophobic face and/or salt bridges of the ApoA-I double belt in HDL particles result in an increase in solvent exposure as well as in flexibility of the protein. It is reasonable to assume that this will affect the lipid dynamics and, hence, both the binding to and the ability to mediate cholesterol efflux. In a recent paper, Manthei et.al.^29^ present a convincing model that indicate the h 4/6 as the site for LCAT-HDL interaction, a site in close proximity to the dynamic h5/5 region. Thus, it is possible to speculate that the affinity of the interaction between LCAT and HDL might change as a consequence of the increased and expanded flexibility around the 4/6 helices observed in ApoA-I amyloidogenic variants in rHDL.

The significance of the findings is not fully clear but may indicate that structural relaxation of this region improves the ApoA-I function as a cholesterol acceptor.

In conclusion, we have shown that the two amyloidogenic amino acid substitutions L75P and L174S lead to increased local structure flexibility, which in turn affects the cholesterol efflux capacity in a positive manner. The importance of understanding the mechanistic details in cholesterol efflux is of general interest and extends beyond explaining the variant-specific differences. For example, it has been shown that impaired cholesterol efflux capacity, due to low^30^ and dysfunctional^31–34^ HDL-cholesterol, may be an important mediator of immunoactivation^33,34^ and subsequent cardiovascular disease in chronic kidney disease^35,36^. Moreover, HDL subpopulation distribution and particle size has been shown to play a role in coronary heart disease people with or without diabetes^36^. The same study concludes that abnormal particle distribution and particle size may contribute to higher risk of developing coronary heart disease in diabetes patients^36^. Therefore, finding solutions to improve (or restore) HDL’s functionality is of interest. This could potentially be done by introducing destabilizing factors that lead to greater region-specific flexibility in the ApoA-I structure. However, in such endeavors, care should be taken to not increase the amyloidogenic potential of the ApoA-I structure.

## Materials and Methods

### Serum samples from patients carrying ApoA-I amyloidogenic variants

Serum samples from patients with ApoA-I amyloidosis and from unrelated control subjects (11 controls, 11 L75P patients and 4 L174S patients, between 37 to 77 years of age, both female and male subjects) were obtained at the Amyloidosis Research and Treatment at Fondazione IRCCS Policlinico San Matteo. Written informed consent for using biological samples and clinical data for research purposes was obtained according to local Institutional review board guidelines.

### Apolipoprotein B depletion

Serum samples from patients carrying ApoA-I amyloidogenic variants and from control subjects were subjected to ApoB depletion prior to perform cholesterol efflux experiments on macrophages, as previously described^37^. For this, 200 μl of serum was incubated with 80 μl of 20 % polyethylene glycol (PEG) 6000 in 20 mM glycine buffer at pH 7.4, for 20 minutes at 25 °C, with gentle shaking. After incubation, serum samples were centrifuged at 10000 rpm for 20 minutes at 4 °C and supernatants were analyzed by human ApoA-I ELISA (3710-1HP-10, Mabtech Inc.), as well as by denaturant and native western blot (procedure is described in the Supplementary Information file).

### Protein expression and purification

Human ApoA-I proteins, containing a His-tag and tobacco etch virus (TEV) protease recognition site at the N-terminus, were expressed in the bacterial *Escherichia coli* (*E. coli*) BL21(DE3) pLysS strain (Invitrogen, Thermo Fisher Scientific) as described in ^38^. Recombinant proteins were then purified using immobilized metal affinity chromatography (His-Trap-Nickel-chelating columns, GE Healthcare) followed by treatment with TEV protease and a second immobilized metal affinity chromatography step to remove the His-tag. Protein purity was analyzed by SDS-PAGE followed by Coomassie staining, and concentration was determined by using a NanoDrop 2000c spectrophotometer (Thermo Fisher Scientific).

### Preparation of reconstituted HDL

Lyophilized POPC (1-palmitoyl-2-oleoyl-sn-glycero-3-phosphocholine, Avanti Polar Lipids) and cholesterol (FC) (Avanti Polar Lipids) were dissolved in 3:1 chloroform:methanol, and the solvent was evaporated by overnight incubation under a stream of nitrogen gas. POPC and FC were dissolved in PBS, and lipoparticles were generated by using the cholate dialysis method^13^. Details about the experimental procedure can be found in the Supplementary Information file.

### Cholesterol efflux from macrophages

Cholesterol efflux assay was performed as described in ^19^. The detailed procedure is described in the Supplementary Information file.

### Synchrotron Radiation Circular Dichroism

SRCD experiments were performed using a nitrogen-flushed Module-A end-station spectrophotometer, equipped with a 6-cell turret, at B23 Beamline at the Diamond Light Source^39–41^. POPC particles were produced in McIlvaine buffer, pH 7, and analyzed by SRCD at 0.15 mg/ml in a 0.2 mm quartz cuvette. Spectra were acquired at 25 °C in the far-UV range 185-260 nm, with 1 nm wavelength increment. All the spectra were corrected by subtracting the background signal of the buffer. Secondary structure estimations from CD spectra was carried out using the software CD Apps^42^ and applying CONTILLN algorithm with reference data SP 43^43^. The molar ellipticity ([Θ]) was calculated according to the equation described in ^44^.

### Hydrogen–deuterium exchange mass spectrometry (HDX-MS)

8.4 nm POPC:ApoA-I particles containing WT, L75P or L174S protein were analysed by HDX at a concentration of 0.3 mg/ml in PBS, at pH 7.4. Two different particle preparations were analysed and data combined for the final HDX analysis. The detailed procedure is described in the Supplementary Information file.

### Thermal stability analyses

CD spectroscopy measurements were performed on a Jasco J-810 spectropolarimeter equipped with a Jasco CDF-426S Peltier. POPC particles (0.1 mg/mL ApoA-I in particle) were diluted in PBS, loaded into a 1 mm quartz cuvette and CD signal at 220 nm was acquired in the 20-98 °C range, with a 2 °C increment. The estimation of the transition temperature (Tm) was performed by biphasic fitting using GraphPad Prism software.

A PCR thermal cycler (TC-Plus, Techne) was used to denature POPC particles with the same temperature increment and incubation times as in the thermal unfolding at the CD spectrometer. Samples were taken at 20, 65, 85 and 98°C and analyzed by native electrophoretic analysis (2.5 μg of ApoA-I per lane) followed by western blot with anti-human ApoA-I antibodies. The quantification of the species visualized on the gel was performed by using ImageJ software and the amount of each species was plotted as fold change with respect to the total signal.

### Statistical analysis

Data shown are the mean ± SD or ± SEM, as indicated. Analysis was performed by one- or two-way ANOVA, as indicated, using the GraphPad Prism software. Outliers were identified using GraphPad outlier test (alpha= 0.05).

## Abbreviations

ABCA1: ATP binding cassette A1
ABCG1: ATP binding cassette G1
ApoA-I: apolipoprotein A-I
ApoB: apolipoprotein B
CD: circular dichroism
CVD: cardiovascular disease
FC: unesterified free cholesterol
HDL: high-density lipoprotein
HDX: hydrogen-deuterium exchange
LCAT: lecithin-cholesteryl acyl transferase
PBS: phosphate buffer saline
POPC: 1-palmitoyl-2-oleoyl-sn-glycero-3-phosphocholine
rHDL: reconstituted HDL
SRCD: Synchrotron Radiation Circular Dichroism
TEV: tobacco etch virus
Tm: transition temperature
WT: wild-type

## Acknowledgments

This work was supported by grants from the Swedish Research council (K2014-54X-22426-01-3 and 2009-1039, Strategic research area Exodiab), the Vinnova (Sweden’s Innovation Agency 2015-01549), the Krapperup Foundation, the Crafoord foundation, the Carl Tesdorpfs foundation, the Royal Physiographic Society in Lund, the Sten K Johnson’s Foundation, and by the Swedish Foundation for Strategic Research (IRC15-0067). Part of the research leading to these results has been supported by the project CALIPSOplus under the Grant Agreement 730872 from the EU Framework Programme for Research and Innovation HORIZON 2020.

## Contribution

JOL and RDG conceived and designed the study. ON, ML, SE and RDG collected the data. LO provided serum samples from patients and control subjects. ON, SE, RDG and JOL analyzed the data. JOL and RDG drafted the manuscript. All the authors revised the manuscript critically and gave their approval for the final version of the manuscript to be published.

## Supplementary Materials and Methods

### Western blot from Native and denaturant gel electrophoresis

Serum samples were separated on 4-15% Tris-Glycine pre-casted gels (BioRad), for the denaturant PAGE and on NativePAGE Bis-Tris Gel System 4-16 % (Invitrogen, Thermo Fisher Scientific), for the native PAGE, according to the manufacturer’s instructions.

In both cases, serum proteins were transferred from the gel to PVDF membranes and probed with anti-human ApoA-I antibodies (64308, Abcam, for denaturant blot and Q0496, Dako, Agilent Technologies, for native blot). Detection was performed by using HRP-conjugated secondary antibodies (GE Healthcare) and a chemiluminescence detection substrate (Super-Signal West Femto, Thermo Fisher Scientific). Blots were imaged using the Odyssey Fc system (LI-COR Biosciences).

### Preparation of reconstituted HDL

POPC and POPC:FC lipoparticles were produced by incubating POPC and cholesterol, diluted in sodium deoxycholate, with ApoA-I variants at 80:4:1 or 40:2:1 molar ratio (to produce 9.6 and 8.4 nm particles, respectively) and at a 1 mg/ml protein concentration. Mixtures were incubated at 37 °C for 1 h and then dialyzed against PBS for 72 h. At the end of the incubation, homogeneous 9.6 nm and 8.4 nm POPC: and POPC:FC:ApoA-I particles were isolated by a size-exclusion chromatography by using a preparative Superose 6 increase 10/300 GL column (GE Healthcare). Samples were eluted at a flow rate of 0.5 ml/min, in PBS, and analyzed by Blue Native PAGE using the NativePAGE Bis-Tris Gel System 4-16 % (Invitrogen, Thermo Fisher Scientific) according to the manufacturer’s instructions, flushed with nitrogen and stored at −80 °C prior to experiments.

### Cholesterol efflux from macrophages

J774 macrophages (TIB-67, ATCC) were plated into 24-well plates, in RPMI 1640 (Gibco) supplemented with 10% FBS and 50 μg/ml gentamicin, at a cell density of 150,000 cells/well. 24 h after the plating, cells were loaded with 4 μCi/ml ^3^H-cholesterol (Perkin Elmer) in RPMI 1640 containing 5% FBS, 2 μg/ml ACAT inhibitor (inhibits formation of cholesteryl esters) (Sandoz 58-035, Sigma) and gentamicin. Upon 24 h incubation, the medium was replaced with RPMI 1640 supplemented with 0.2% BSA (low free fatty acids and low endotoxin, Sigma), 2 μg/ml ACAT inhibitor, 0.3 mM Cpt-cAMP (promotes expression of ABCA1) (Abcam) and gentamicin for 18 h. At the end of incubation, cells were washed twice with serum free RPMI 1640 and then triplicate wells were treated with either ApoA-I amyloidogenic variants or the WT protein in POPC particles, or serum from patients and control subjects, in RPMI 1640 supplemented with 0.2% BSA, at the indicated concentrations and for 4 h. Cholesterol efflux was measured by collecting the media, centrifuging at 4000 × g for 5 min at room temperature, and transferring of 100 μl supernatant to a scintillation vial. 5 ml of scintillation fluid was added to each sample before scintillation counting was performed. The measure of the total cellular ^3^H-cholesterol was obtained by incubating the cells, in triplicate, with 1% sodium deoxycholate and lysates collected for scintillation counting. Efflux for each treatment was calculated as % of the total ^3^H-cholesterol. Spontaneous basal efflux was measured in triplicate and the efflux for each treatment subtracted for this value.

### Hydrogen–deuterium exchange mass spectrometry (HDX-MS)

All chemicals for the HDX-MS analyses were purchased from Sigma Aldrich, except n-Dodecyl-b-D-Maltopyranoside (DDM) which was from Thermo Scientific. pH measurements were made using a SevenCompact pH-meter equipped with an InLab Micro electrode (Mettler-Toledo) and, prior to all measurements, a 4-point calibration (pH 2,4,7,10) was performed.

The ApoA-I particles were placed in the autosampler in such a way that no sample had a permanence in the machine for longer than 12 hours.

The HDX-MS analysis was performed using automated sample preparation on a LEAP H/D-X PAL™ platform interfaced to an LC-MS system, comprising an Ultimate 3000 micro-LC coupled to an Orbitrap Q Exactive Plus MS. For the HDX-MS, 5 μl of POPC:ApoA-I particles were diluted either in 25 μl of PBS, pH 7.4 or in HDX labelling buffer (PBS prepared in D_2_O, pH_(read)_ 7.0) and the HDX reactions were carried out for t = 0, 30, 300, 3000 and 9000 s at 8°C. At the end of incubation, labelling was quenched by adding 25 μl of 1 % TFA, 0.2 % DDM, 4 M urea, pH 2.5 at 1°C, to the samples. Then, 50 μl of the quenched sample were directly subjected to online pepsin digestion at 4 °C, by injection on a pepsin column (2.1 × 30 mm, Life Technologies). In order to remove lipids from the samples, the pepsin column was directly followed by a 2 × 20 mm guard column (Upchurch Scientific) packed with a washed and equilibrated ZrO_2_ material (Sigma Aldrich, Zirconium IV oxide, powder, <5 micron). The online digestion and trapping were performed for 4 minutes using a flow of 50 μL/min 0.1 % formic acid (FA), pH 2.5. Peptides generated by pepsin digestion were subjected to on-line SPE on a PepMap300 C18 trap column (1 mm × 15 mm) and washed with 0.1% FA for 60 s. Thereafter, the trap column was switched in-line with a reversed-phase analytical column, Hypersil GOLD, particle size 1.9 μm, 1 × 50 mm, and separation was performed at 1°C using a gradient of 5-50 % B over 8 minutes and then from 50 to 90 % B for 5 minutes, the mobile phases were 0.1 % FA (A) and 95 % acetonitrile/0.1 % FA (B).

Following separation, the trap and column were equilibrated at 5 % organic content, until the next injection. The needle port and sample loop were cleaned three times after each injection with mobile phase 5 % MeOH/0.1 % FA, followed by 90 % MeOH/0.1 % FA and a final wash of 5 % MeOH/0.1 % FA. After each sample and blank injection, the pepsin column was washed by injecting 90 μl of pepsin wash solution 1 % FA /4 M urea /5 % MeOH. In order to minimize carry-over, a full blank was run between each sample injection. Separated peptides were analysed on a Q Exactive Plus MS, equipped with a HESI source operated at a capillary temperature of 250 °C. For the undeuterated samples (t = 0 s), injections were acquired using data dependent MS/MS HCD for identification of generated peptides. For the HDX analysis (all labelled samples and one t= 0 s), MS full scan spectra at a setting of 70 K resolution, AGC 3e6, Max IT 200 ms and scan range 300-2000 were collected.

PEAKS Studio 8.5 Bioinformatics Solutions Inc. (BSI, Waterloo, Canada) was used for peptide identification after pepsin digestion of undeuterated samples (i.e. 0 s time point). The search was done on a FASTA file with only the three different ApoA-I sequences; search criteria included a mass error tolerance of 15 ppm and a fragment mass error tolerance of 0.05 Da, oxidation of methionine (15.99 Da) as variable modification and allowing for fully unspecific cleavage by pepsin.

Peptides identified by PEAKS with a peptide score value of log P > 25 and no oxidation were used to generate a peptide list containing peptide sequences, charge state and retention time for the HDX analysis. HDX data analysis and visualization was performed using HDExaminer, version 2.5 (Sierra Analytics Inc., Modesto, US). Due to the comparative nature of the measurements, the deuterium incorporation levels for the peptic peptides were derived from the observed mass difference between the deuterated and non-deuterated peptides without back-exchange correction using a fully deuterated sample. HDX data was normalized to 100 % D_2_O content with an estimated average deuterium recovery of 75 %.

The peptide deuteration of a peptide is the average of all high and medium confidence results and the two first residues assumed unable to hold deuteration. The allowed retention time window was ± 0.5 minute. Heatmaps settings were uncoloured proline, heavy smoothing and the difference heatmaps were drawn using the residual plot as significance criterion (± 0.5 Da). Since previous studies have described bimodal HDX kinetics for regions of ApoA-I in HDL particles^1^, data was analysed allowing the software to try an EX1 deuteration envelope if the EX2 score was lower than 0.9, and to accept the result if the score increased ≥ 0.05. The spectra for all timepoints were manually inspected; low scoring peptides, outliers and peptides were retention time correction could not be made consistent were removed.

## Supplementary Figure Legends

**Figure S1.**
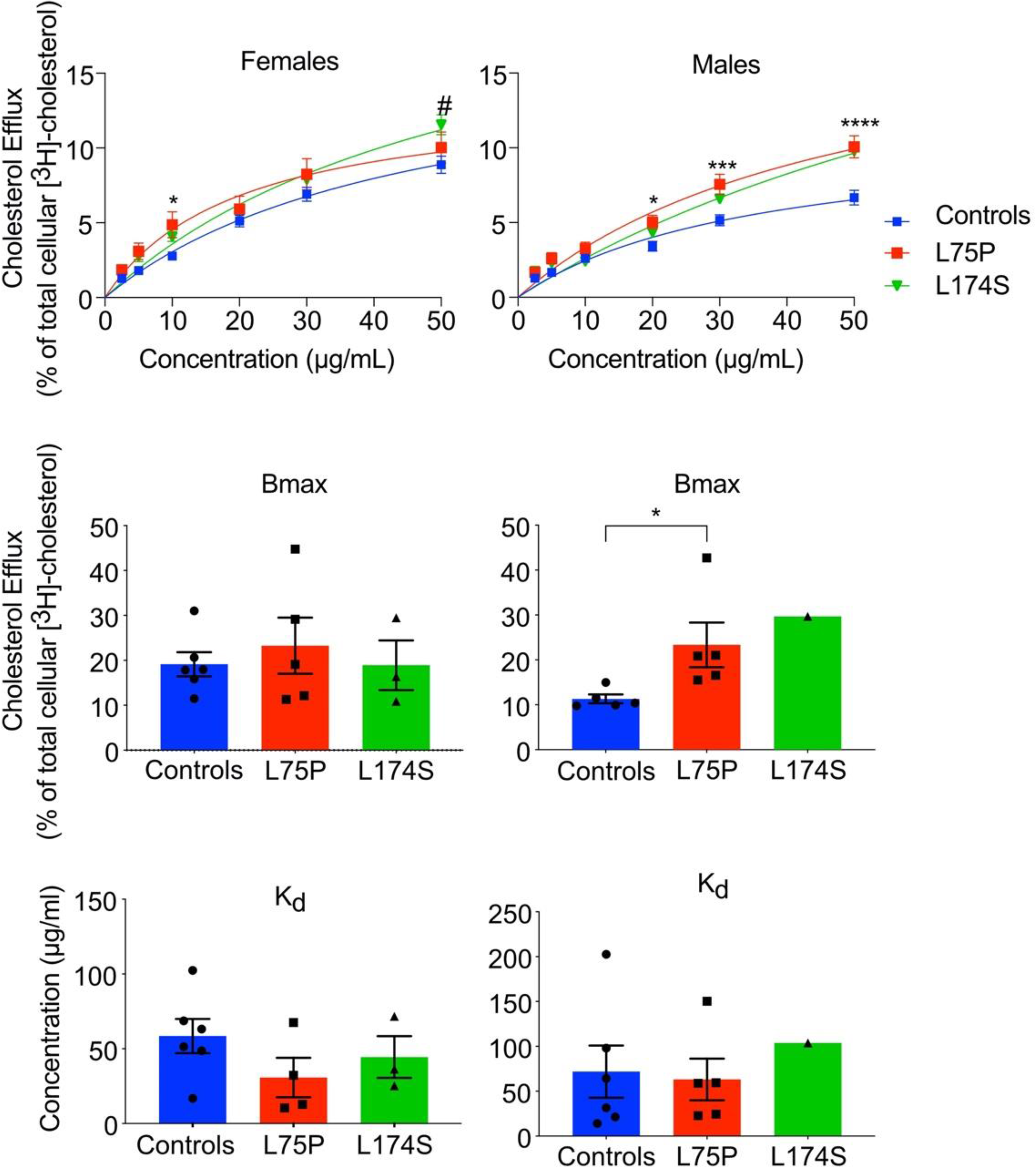
The improved efflux capacity of the amyloidogenic ApoA-I patients is sex specific. Cholesterol efflux data obtained from experiment shown in Figure 1c were plotted according to donor’s sex. Experimental data (upper panels), calculated Bmax (middle panels) and Kd (lower panels). Data shown are the mean ± SEM and significance is calculated according to two-way ANOVA (*p<0.05, ***p<0.001, ****p<0.0001 for L75P patients as compared to healthy donors).

**Figure S2.**
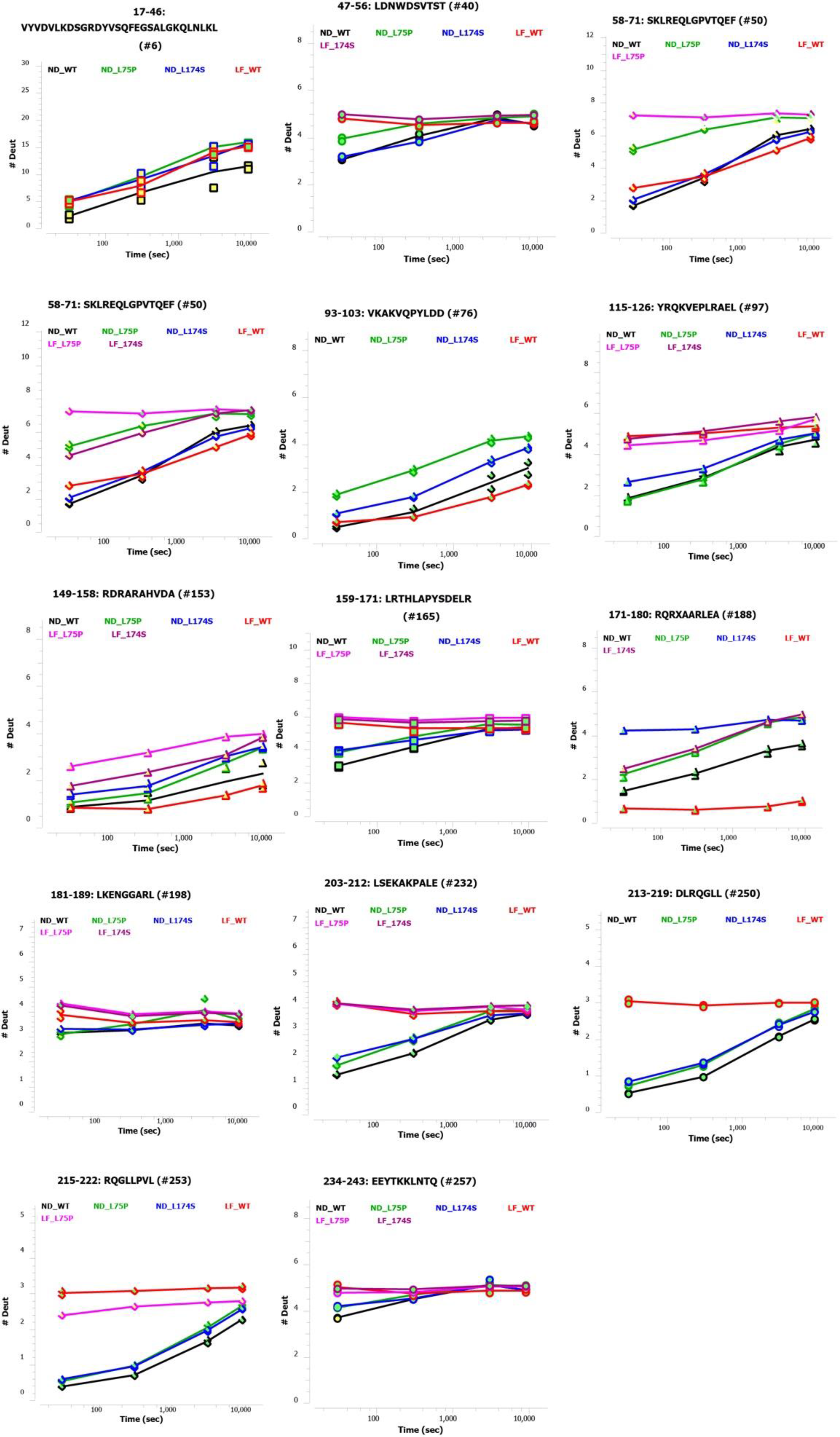
Deuterium uptake of selected ApoA-I peptides. Individual deuterium uptake plots for ApoA-I in 8.4 nm POPC particles (nanodiscs, ND), or in the lipid free (LF) form. LF data is from a single measurement. The peptide sequence is reported above each individual plot. Black trace (ND_WT), red (LF_WT), green (ND_L75P), magenta (LF_L75P), blue (ND_L174S) and pink (LF_L174S).

**Figure S3.**
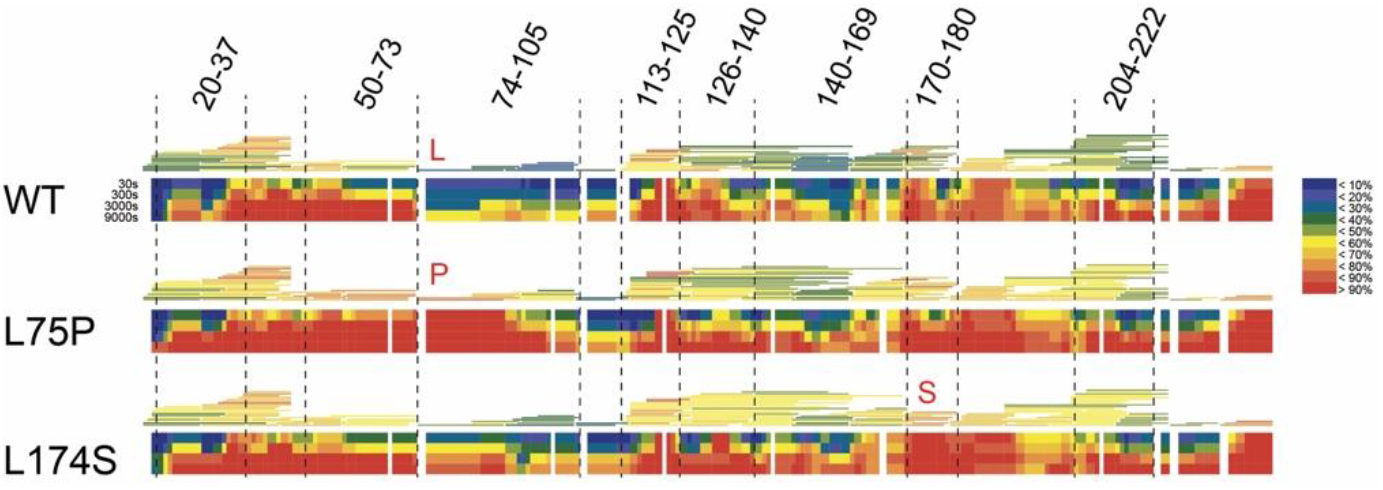
Individual HDX heatmaps of WT, L75P and L174S ApoA-I in 8.4 nm rHDL. HDX peptide coverage is shown by the bars above each heatmap, color-coded to the average deuterium uptake over the 4 observed time points (30 s, 300 s, 3000 s and 9000 s). The color coding is based on a least square calculation of observed deuterium uptake with moderate smoothing. Cold color = slower exchange; warm color = faster exchange. see color key to the right. Individual deuterium uptake curves for all observed peptides can be found in the supplementary file, Supplementary data file S1.pdf.

## References

1. Chiti, F. & Dobson, C.M. Protein Misfolding, Amyloid Formation, and Human Disease: A Summary of Progress Over the Last Decade. Annu Rev Biochem 86, 27–68 (2017).

2. Favari, E., et al. Cholesterol efflux and reverse cholesterol transport. Handb Exp Pharmacol 224, 181–206 (2015).

3. Zannis, V.I., Chroni, A. & Krieger, M. Role of apoA-I, ABCA1, LCAT, and SR-BI in the biogenesis of HDL. J Mol Med (Berl) 84, 276–294 (2006).

4. Phillips, M.C. Molecular mechanisms of cellular cholesterol efflux. J Biol Chem 289, 24020–24029 (2014).

5. Lagerstedt, J.O., et al. Structure of apolipoprotein A-I N terminus on nascent high density lipoproteins. J Biol Chem 286, 2966–2975 (2011).

6. Du, X.M., et al. HDL particle size is a critical determinant of ABCA1-mediated macrophage cellular cholesterol export. Circ Res 116, 1133–1142 (2015).

7. Rye, K.A. & Barter, P.J. Cardioprotective functions of HDLs. J Lipid Res 55, 168–179 (2014).

8. Obici, L., et al. Structure, function and amyloidogenic propensity of apolipoprotein A-I. Amyloid 13, 191–205 (2006).

9. Rowczenio, D., et al. Amyloidogenicity and clinical phenotype associated with five novel mutations in apolipoprotein A-I. Am J Pathol 179, 1978–1987 (2011).

10. Ramella, N.A., et al. Human apolipoprotein A-I natural variants: molecular mechanisms underlying amyloidogenic propensity. PLoS One 7, e43755 (2012).

11. Gursky, O., Mei, X. & Atkinson, D. The crystal structure of the C-terminal truncated apolipoprotein A-I sheds new light on amyloid formation by the N-terminal fragment. Biochemistry 51, 10–18 (2012).

12. Arciello, A., Piccoli, R. & Monti, D.M. Apolipoprotein A-I: the dual face of a protein. FEBS Lett 590, 4171–4179 (2016).

13. Ray, A., et al. Specific Cholesterol Binding Drives Drastic Structural Alterations in Apolipoprotein A1. J Phys Chem Lett 9, 6060–6065 (2018).

14. Rader, D.J., et al. In vivo metabolism of a mutant apolipoprotein, apoA-IIowa, associated with hypoalphalipoproteinemia and hereditary systemic amyloidosis. J Lipid Res 33, 755–763 (1992).

15. Obici, L., et al. Liver biopsy discloses a new apolipoprotein A-I hereditary amyloidosis in several unrelated Italian families. Gastroenterology 126, 1416–1422 (2004).

16. Gomaraschi, M., et al. Effect of the amyloidogenic L75P apolipoprotein A-I variant on HDL subpopulations. Clin Chim Acta 412, 1262–1265 (2011).

17. Obici, L., et al. The new apolipoprotein A-I variant leu(174) --> Ser causes hereditary cardiac amyloidosis, and the amyloid fibrils are constituted by the 93-residue N-terminal polypeptide. Am J Pathol 155, 695–702 (1999).

18. Muiesan, M.L., et al. Vascular alterations in apolipoprotein A-I amyloidosis (Leu75Pro). A case-control study. Amyloid 22, 187–193 (2015).

19. Del Giudice, R., et al. Structural determinants in ApoA-I amyloidogenic variants explain improved cholesterol metabolism despite low HDL levels. Biochim Biophys Acta Mol Basis Dis 1863, 3038–3048 (2017).

20. Gregorini, G., et al. Tubulointerstitial nephritis is a dominant feature of hereditary apolipoprotein A-I amyloidosis. Kidney Int 87, 1223–1229 (2015).

21. Diditchenko, S., et al. Novel formulation of a reconstituted high-density lipoprotein (CSL112) dramatically enhances ABCA1-dependent cholesterol efflux. Arterioscler Thromb Vasc Biol 33, 2202–2211 (2013).

22. Segrest, J.P., et al. The amphipathic helix in the exchangeable apolipoproteins: a review of secondary structure and function. J Lipid Res 33, 141–166 (1992).

23. Segrest, J.P., et al. A detailed molecular belt model for apolipoprotein A-I in discoidal high density lipoprotein. J Biol Chem 274, 31755–31758 (1999).

24. Chetty, P.S., et al. Comparison of apoA-I helical structure and stability in discoidal and spherical HDL particles by HX and mass spectrometry. J Lipid Res 54, 1589–1597 (2013).

25. Sevugan Chetty, P., et al. Apolipoprotein A-I helical structure and stability in discoidal high-density lipoprotein (HDL) particles by hydrogen exchange and mass spectrometry. Proc Natl Acad Sci U S A 109, 11687–11692 (2012).

26. Morgan, C.R., et al. Conformational transitions in the membrane scaffold protein of phospholipid bilayer nanodiscs. Mol Cell Proteomics 10, M111 010876 (2011).

27. Wilson, C.J., Das, M., Jayaraman, S., Gursky, O. & Engen, J.R. Effects of Disease-Causing Mutations on the Conformation of Human Apolipoprotein A-I in Model Lipoproteins. Biochemistry 57, 4583–4596 (2018).

28. Hudgens, J.W., et al. Interlaboratory Comparison of Hydrogen-Deuterium Exchange Mass Spectrometry Measurements of the Fab Fragment of NISTmAb. Anal Chem 91, 7336–7345 (2019).

29. Manthei, K.A., et al. Structural analysis of lecithin:cholesterol acyltransferase bound to high density lipoprotein particles. Commun Biol 3, 28 (2020).

30. Muntner, P., He, J., Astor, B.C., Folsom, A.R. & Coresh, J. Traditional and nontraditional risk factors predict coronary heart disease in chronic kidney disease: results from the atherosclerosis risk in communities study. J Am Soc Nephrol 16, 529–538 (2005).

31. Holzer, M., et al. Uremia alters HDL composition and function. J Am Soc Nephrol 22, 1631–1641 (2011).

32. Yamamoto, S., et al. Dysfunctional high-density lipoprotein in patients on chronic hemodialysis. J Am Coll Cardiol 60, 2372–2379 (2012).

33. Speer, T., et al. Abnormal high-density lipoprotein induces endothelial dysfunction via activation of Toll-like receptor-2. Immunity 38, 754–768 (2013).

34. Weichhart, T., et al. Serum amyloid A in uremic HDL promotes inflammation. J Am Soc Nephrol 23, 934–947 (2012).

35. Rogacev, K.S., et al. Lower Apo A-I and lower HDL-C levels are associated with higher intermediate CD14++CD16+ monocyte counts that predict cardiovascular events in chronic kidney disease. Arterioscler Thromb Vasc Biol 34, 2120–2127 (2014).

36. Tian, L., et al. High-density lipoprotein subclass and particle size in coronary heart disease patients with or without diabetes. Lipids Health Dis 11, 54 (2012).

37. Davidson, W.S., et al. The effects of apolipoprotein B depletion on HDL subspecies composition and function. J Lipid Res 57, 674–686 (2016).

38. Del Giudice, R. & Lagerstedt, J.O. High-efficient bacterial production of human ApoA-I amyloidogenic variants. Protein Sci 27, 2101–2109 (2018).

39. Siligardi, G. & Hussain, R. CD spectroscopy: an essential tool for quality control of protein folding. Methods Mol Biol 1261, 255–276 (2015).

40. Hussain, R., Javorfi, T. & Siligardi, G. Circular dichroism beamline B23 at the Diamond Light Source. J Synchrotron Radiat 19, 132–135 (2012).

41. Javorfi, T., Hussain, R., Myatt, D. & Siligardi, G. Measuring circular dichroism in a capillary cell using the b23 synchrotron radiation CD beamline at diamond light source. Chirality 22 Suppl 1, E149–153 (2010).

42. Hussain, R., et al. CDApps: integrated software for experimental planning and data processing at beamline B23, Diamond Light Source. Corrigendum. J Synchrotron Radiat 22, 862 (2015).

43. Provencher, S.W. & Glockner, J. Estimation of globular protein secondary structure from circular dichroism. Biochemistry 20, 33–37 (1981).

44. Brouillette, C.G., et al. Forster resonance energy transfer measurements are consistent with a helical bundle model for lipid-free apolipoprotein A-I. Biochemistry 44, 16413–16425 (2005).

## Supplementary References

1. Wilson, C.J., Das, M., Jayaraman, S., Gursky, O. & Engen, J.R. Effects of Disease-Causing Mutations on the Conformation of Human Apolipoprotein A-I in Model Lipoproteins. Biochemistry 57, 4583–4596 (2018).

